# Serotonin coordinates reproductive functions in *Caenorhabditis elegans*

**DOI:** 10.1101/2022.05.11.491580

**Authors:** Erin Z. Aprison, Svetlana Dzitoyeva, Ilya Ruvinsky

**Affiliations:** Department of Molecular Biosciences, Northwestern University, Evanston, IL 60208, USA

**Keywords:** Serotonin, *C. elegans*, pheromone, germline, reproduction, coordination

## Abstract

Reproduction alters animal behavior and physiology, but neuronal circuits that coordinate these changes remain largely unknown. Insights into mechanisms that regulate and possibly coordinate reproduction-related traits could be gleaned from the study of sex pheromones that manipulate potential mating partners to improve reproductive success. In *C. elegans*, the prominent male pheromone, ascr#10, modifies reproductive behavior and several aspects of reproductive physiology in hermaphrodite recipients, including improving oocyte quality. Here we show that a circuit that contains serotonin-producing and serotonin-uptaking neurons plays a key role in mediating these beneficial effects of ascr#10. We also demonstrate that increased serotonergic signaling promotes proliferation of germline progenitors in adult hermaphrodites. Our results establish a role for serotonin in maintaining germline quality and highlight a simple neuronal circuit that acts as a linchpin that couples food intake, mating behavior, reproductive output, and germline renewal and provisioning.

## INTRODUCTION

Males and females of the same species employ a rich repertoire of signals to improve reproductive success. Sex pheromones that modulate behavior and reproductive physiology of potential mates are a prominent tool in this arsenal of manipulation (1). The case of the most abundant male-biased ascaroside pheromone in *C. elegans*, ascr#10 (2), is instructive. This small molecule alters multiple behaviors. Hermaphrodites exposed to physiological concentrations of ascr#10 reduce exploratory movement (3). A shift from global to local exploration, respectively referred to as roaming and dwelling (4–6), is generally associated with increased exploitation of local resources (7). ascr#10 also increases mating receptivity in hermaphrodites and promotes egg-laying in already reproducing animals (3).

In addition to these behavioral changes, ascr#10 affects several aspects of hermaphrodite reproductive physiology. It improves sperm guidance (8) and increases stores of germline precursor cells (GPCs) in older worms (9). Exposure to ascr#10 improves quality of the oogenic germline (10). This manifests in a more youthful oocyte morphology, decreased rates of chromosomal nondisjunction, and lower embryonic lethality both in the wild-type and mutant genetic backgrounds (10). A likely mechanism responsible for the improved oocyte quality on ascr#10 begins with greater mitotic proliferation in the germline. The more numerous germline precursor cells undergo physiological cell death thus scavenging resources (nutrients, metabolites, organelles, etc.) that could be used to improve the quality of the few surviving oocytes (10). That an external signal can elicit these effects demonstrates that the nervous system can regulate germline quality, but the specific circuits are not known.

Some effects of ascr#10 are known to require the function of a specific serotonergic circuit that contains NSM and HSN neurons that signal via the *mod-1* receptor (3, 11), the same circuit that regulates the division of locomotor behavior into episodes of roaming and dwelling (6). We therefore tested whether the same serotonergic signaling regulates quality of the oogenic germline.

## RESULTS AND DISCUSSION

### The serotonergic circuit that is required for ascr#10 improvement of oocyte quality

Of the five classes of serotonergic neurons in *C. elegans* hermaphrodites, three (NSM, ADF, and HSN) express a serotonin biosynthetic enzyme TPH-1 and can therefore produce serotonin, whereas two (AIM and RIH) can only uptake serotonin synthesized elsewhere (12). Increased serotonergic signaling from two classes of producing neurons, NSM and HSN, acting via the MOD-1 receptor mediates effects of ascr#10 on exploratory behavior and GPC counts in older hermaphrodites (3, 13). We sought to test whether the NSM/HSN/MOD-1 neuronal circuit also mediates the salubrious effects of ascr#10 on the quality of the oogenic germline. An expedient assay of quality is whether an oocyte could support successful embryonic development. In wild-type N2 hermaphrodites that were mated just after exhausting their self-sperm supply, exposure to ascr#10 nearly halved embryonic lethality; loss-of-function alleles of *tph-1* or *mod-1* precluded this quality improvement (Figure 1A). A decrease in embryonic lethality was also observed in self-broods of hermaphrodites maintained in the presence of ascr#10, but the effect was not observed in *tph-1* and *mod-1* mutants (Figure 1B). The ascr#10 pheromone improves oocyte quality in older mothers because it stimulates proliferation of germline precursors and physiological cell death in the adult germline (10). In the absence of serotonergic signaling, ascr#10 did not increase either proliferation or cell death in the germline (Figure 1C, D).

**Figure 1.**
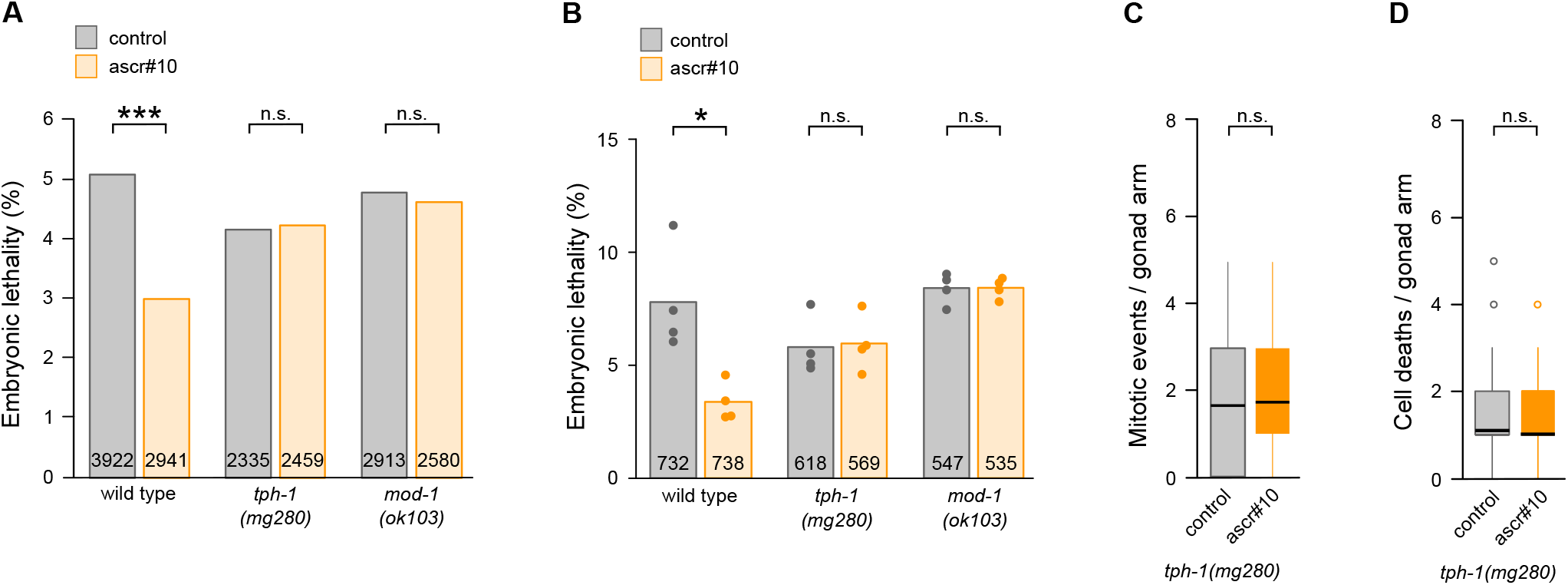
Serotonergic signaling is required for the salubrious effects of ascr#10 on the oogenic germline. **(A)** Percentage of unhatched embryos on vs off ascr#10 in the progeny of N2, *tph-1*, and *mod-1* self-sperm depleted hermaphrodites mated on Day 5 of adulthood to young males. Total numbers of tested embryos are indicated inside relevant bars. **(B)** Percentage of unhatched embryos on vs off ascr#10 in the self-progeny of N2, *tph-1*, and *mod-1* hermaphrodites during Days 4 & 5 of adulthood. Total numbers of tested embryos are indicated inside relevant bars. Dots represent percentage of embryonic lethality in independent experiments. **(C)** Unlike wild-type N2 hermaphrodites (10), *tph-1* hermaphrodites show no increase of germline precursor divisions in the presence of ascr#10. Quantified using phospho-Histone 3 (pH3) staining during Day 2 of adulthood. **(D)** Unlike wild-type N2 hermaphrodites (10), *tph-1* hermaphrodites show no increase of germline cell death in the presence of ascr#10. Quantified using SYTO12 staining during Day 3 of adulthood. Black bars denote means. Asterisks indicate levels of statistical significance (* for p<0.05, *** for p<0.001). Binomial test in **A** and **B**, Kolmogorov-Smirnov test in **C** and **D**. See Table S1 for primary data and details of statistical analyses.

### Increased serotonin signaling promotes germline proliferation

Loss of function mutations in the serotonin transporter gene *mod-5* reduce serotonin reuptake, effectively increasing serotonergic signaling (14). The number of GPCs in the otherwise untreated hermaphrodites carrying *mod-5(lf)* mutations was significantly higher than in any of the 31 other mutant strains we tested (Figure 2A), suggesting that higher levels of serotonin signaling lead to the increase in the number of GPCs. The *mod-5* gene is expressed in four classes of serotonergic neurons including producing (NSM and ADF) and uptaking-only (AIM and RIH) cells (12, 15). Whereas expressing MOD-5 in the producing NSM and ADF neurons did not rescue the *mod-5* defect, expression in a set of cells that included AIMs and RIH, but no serotonin-producing neurons, restored the wild-type GPC numbers (Figure 2A).

**Figure 2.**
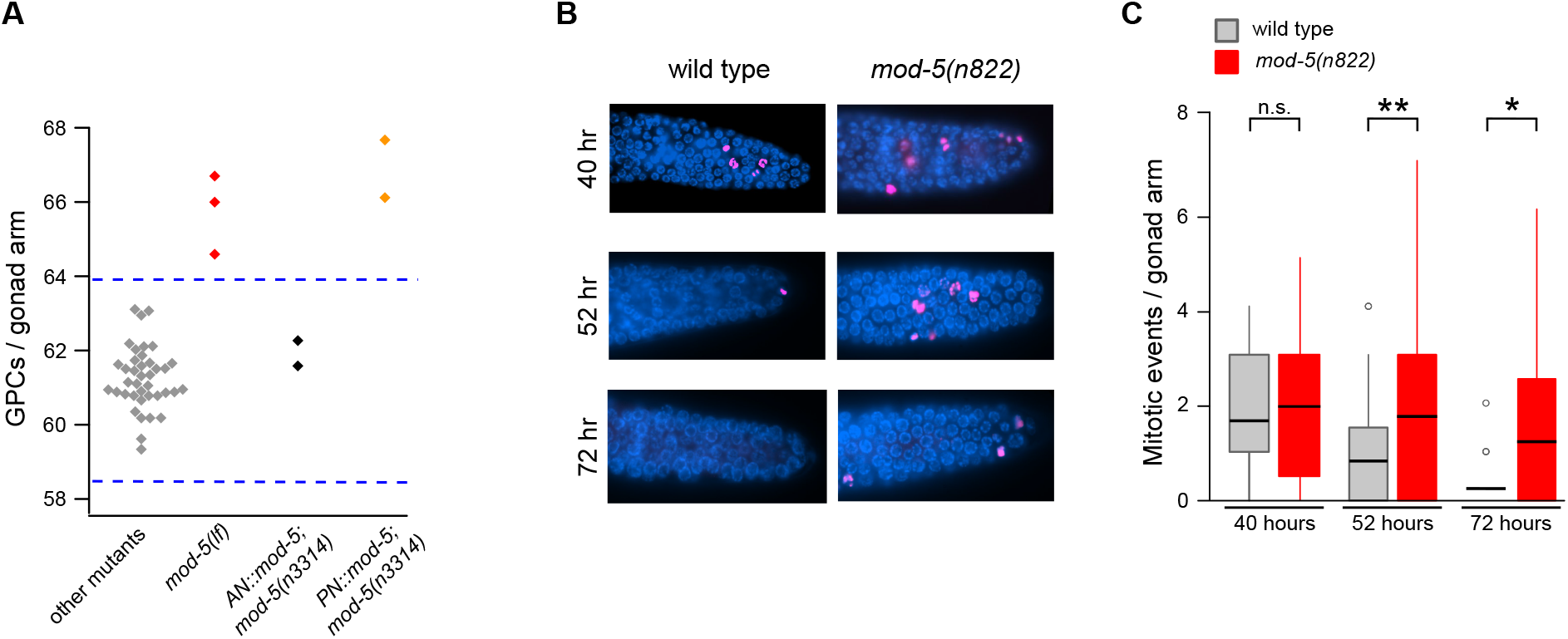
Increased serotonergic signaling increases the number of germline precursors. **(A)** Comparison of GPC counts from 39 experiments using 31 mutants tested in (3), three experiments involving *mod-5* hermaphrodites [two experiments using *mod-5(n822)* and one experiment using *mod-5(n3314)]*, two experiments using the strain that restores MOD-5 function in neurons including AIM and RIH (*AN::mod-5*), and two experiments using the strain that restores MOD-5 function in neurons including NSM and ADF (*PN::mod-5*). Each diamond represents the mean value from one experiment. Dashed lines delimit three standard deviations above and below the mean of all strains except *mod-5*. See Figure S1 for more detail. **(B)** Representative images of pH3 staining of gonads in wild-type N2 and *mod-5* hermaphrodites aged mid-L4 (40 hours), pre-reproductive adult (52 hours), and Day 2 of adulthood (72 hours). **(C)** Quantification of cell divisions (pH3 staining) in the Progenitor Zone in the germlines of N2 and *mod-5* hermaphrodites. In none of the experiments in this figure, were hermaphrodites treated with ascr#10. Asterisks indicate levels of statistical significance (* for p<0.05, ** for p<0.01). Kolmogorov-Smirnov test in **C**. See Table S1 for primary data and details of statistical analyses.

Proliferation of the germline precursor cells is reduced in adult *C. elegans* hermaphrodites compared to late larvae (16). Exposure to ascr#10 maintained higher precursor proliferation in wild-type adult hermaphrodites (10). The untreated *mod-5(lf)* adults hermaphrodites showed increased germline proliferation, comparable to that of ascr#10-exposed wild-type worms (Figure 2B, C).

### A model for coordinated effects of serotonin on *C. elegans* reproduction

Our results implicate the majority of serotonergic neurons in *C. elegans* hermaphrodites in mediating reproductive functions (Figure 3). Exposure to ascr#10 increases serotonin signaling from NSM and HSN neurons acting via the MOD-1 receptor (3, 13). The pharyngeal NSM neurons release serotonin upon food ingestion (17). Consistently, ascr#10 promotes consumption of local resources at the expense of exploration (3). Pheromone-reduced locomotion, either alone or in combination with other factors, increases the likelihood of successful mating (3).

**Figure 3.**
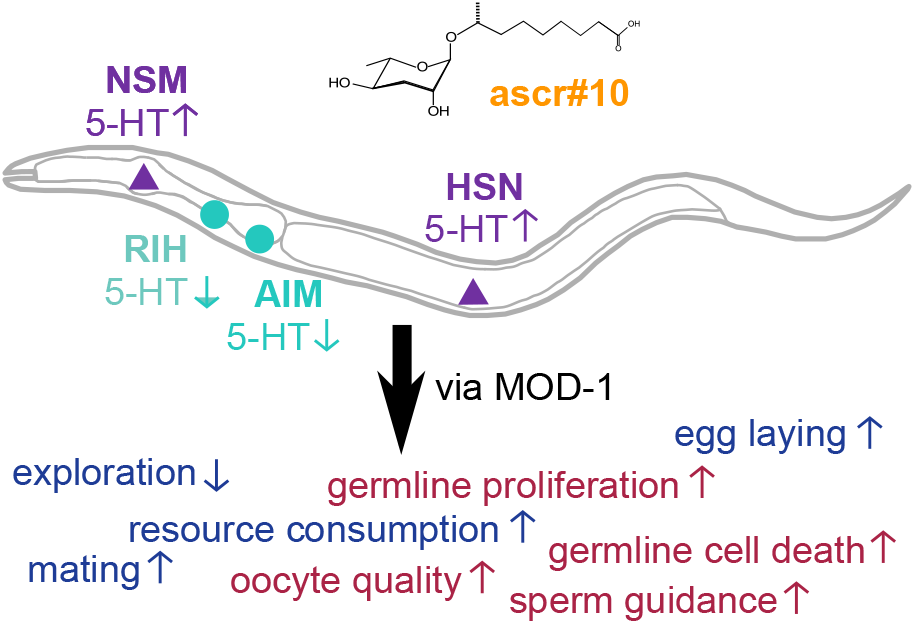
The serotonergic circuit that coordinately regulates multiple reproductive functions in *C. elegans*. A hypothesis regarding the role of serotonin in promoting reproductive behavior and physiology. Observed reproduction-related effects can be loosely categorized as behavioral (blue) or physiological (red). See text for details.

Serotonin release from HSN neurons stimulates egg laying (18) as does exposure to ascr#10 (3). This male pheromone improves sperm guidance in the hermaphrodite reproductive tract (8); sperm guidance is positively regulated by serotonergic signaling (19). Notably, ascr#10 increases germline proliferation in hermaphrodites (10). Here we showed that serotonergic signaling is required for this effect, while serotonin uptake via MOD-5 (acting in AIM and RIH) reduces germline proliferation. We conclude that increased serotonergic signaling increases production of germline precursors, presumably to replenish oocyte stores depleted by greater egg laying (i.e., offspring production) in the presence of the pheromone. In addition to increasing the number of germline cells in hermaphrodites, ascr#10 also increases their quality (10).

Therefore, a major role of the circuit that consists of serotonin-producing NSM and HSN neurons, the MOD-1 receptor, and the MOD-5 transporter in AIM and RIH neurons in *C. elegans* hermaphrodites is to coordinate resource consumption required for greater reproductive output, with reproduction-promoting behaviors, and to match quality and quantity of produced oocytes to demands imposed by germline expenditure and the presence of potential mates. The male pheromone ascr#10 stimulates activity of this circuit to manipulate hermaphrodites to direct greater resources to progeny production.

### Is serotonin a conserved regulator of reproduction?

There is considerable evidence that serotonin regulates various reproduction-related traits in different species and at least some aspects of the regulatory logic we described here may be broadly conserved. For example, serotonergic signaling reduces exploratory movement in *C. elegans* (3, 6), *D. melanogaster* (20), and mice (21). In *D. melanogaster*, serotonin is involved in regulating a dietary switch that occurs in females following mating and manages the balance of nutrients, a role that may be conserved with other animals (22). Serotonin has also been implicated in a variety of reproductive processes. In Drosophila females, mating causes changes in the levels and distribution of serotonin in the termini of neurons that innervate reproductive organs (23). In mammals, the effects of serotonin on reproduction-related behaviors (24) and on the germline (25) are varied and complex. Some, but not all, roles played by serotonin in regulating behavioral and physiological aspects of reproduction in mammals may be shared with other vertebrates (26). Likewise, insects have shown diverse responses to increases in serotonin – while in mosquitoes serotonin was seen as promoting ovarian development (27), in Drosophila increased serotonergic signaling caused oogenesis defects (28). It remains to be determined whether the apparent differences are due to divergence in the underlying mechanisms or to discrepancies in the experimental set up. Environmental conditions may be one particularly saliant variable – for example, in Drosophila, serotonin enhances ovarian dormancy, an arrest of female gonadal maturation, under adverse circumstances (29). Future comparative work will benefit from standardizing experimental paradigms across species. We also suggest that precise definition of the circuits (sources of signal, receptors, sites of action) involved in regulating specific behaviors and physiological processes, as we reported here, will be particularly important for inferring conserved roles of serotonin in regulating reproductive functions in animals.

## MATERIALS AND METHODS

The following stains were used: wild type N2 (CGC), MT15434 *tph-1(mg280)* (CGC), MT9668 *mod-1(ok103)* (CGC), AN::*mod-5 mbr-1p::mod-5;mod-5(n3314)* (Sze lab), *PN::mod-5 tph-1p::mod-5;mod-5(n3314)* (Sze lab), MT8944 *mod-5(n822)* (Koelle lab), MT9772 *mod-5(n3314)* (CGC). Standard, previously published methods were used (3, 10, 11, 13, 30). All experimental treatments were processed in parallel with matched controls. Worms were synchronized by hypochlorite treatment and overnight incubation in M9, after which the synchronized L1 larvae were placed (in small populations, 10 or 30 worms, depending on the experiment) on agar plates seeded with *E. coli* OP50. Pre-reproductive adults (48 hours post release from L1 arrest) were transferred to OP50-seeded plates that were either control or conditioned with synthetic ascr#10 (gift of F. C. Schroeder). For assessing embryonic lethality following mating, Day 5 hermaphrodites (these have exhausted self-sperm) were singled and mated to one young male for 2 hours. Numbers of live and dead progeny were counted for 3 days thereafter. For assessing embryonic lethality in self-broods, progeny produced during the last day of self-fertility (end of Day 4 through Day 5 of adulthood) were considered. More detailed protocol can be found in (10). For quantifying germline proliferation, gonads were dissected and stained with Anti-Histone H3 (phospho S10) antibodies following a modified protocol of (31), as described in (10), and only prophase nuclei were counted. For counting cell death events we used SYTO12 (Invitrogen) and the protocol by (32) as detailed in (10). For GPC counts, on Day 5 of adulthood, hermaphrodites were stained with DAPI (4’,6-diamidino-2-phenylindole) as described (9) and the germline precursor cells were counted. The boundary between the Progenitor Zone and the more proximal transition zone is defined by the appearance of crescent-shaped nuclei that have progressed to leptotene/zygotene stages of meiotic prophase (33, 34). The *AN::mod-5* (“absorbing neurons”) and *PN::mod-5* (“producing neurons”) strains (gift of J. Y. Sze) were reported in (12). Expression of the *AN::mod-5* was directed by a *mbr-1* promoter. Expression of the *mbr-1* gene can be detected in 28 classes of neurons (15), but the only overlap with the pattern of *mod-5* expression is in AIM and RIH neurons. Expression of the *PN::mod-5* was directed by the BCD region of the *tph-1* promoter (35). The overlap between the expression pattern of this promoter and the *mod-5* gene is in two neurons: NSM and ADF.

## Acknowledgments

We thank R. Morimoto for generous hospitality, J.Y. Sze and M. Koelle for strains, and F. Schroeder for ascr#10. We thank WormBase and the Caenorhabditis Genetics Center (CGC). WormBase is supported by grant U41 HG002223 from the National Human Genome Research Institute at the NIH, the UK Medical Research Council, and the UK Biotechnology and Biological Sciences Research Council. The CGC is funded by the NIH Office of Research Infrastructure Programs (P40 OD010440).

