## Supplementary material for "Serotonin coordinates reproductive functions in *Caenorhabditis elegans*": Figure S1

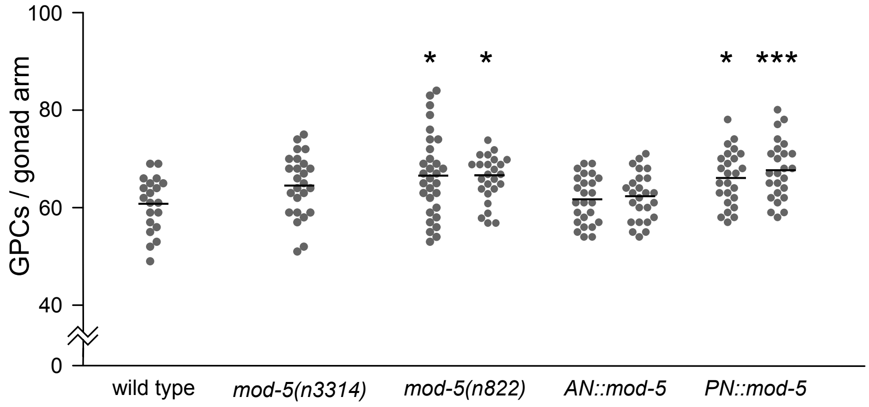

GPCs / gonad arm
100
80
60
40
0
* *
* ***
wild type
mod-5(n3314)
mod-5(n822)
AN::mod-5
PN::mod-5

**Figure S1. Numbers of GPCs in several mutants shown in Figure 2A.**

Number of GPCs per gonad arm in hermaphrodites on Day 5 of adulthood in several strains shown in Figure 2A. Wild type (N2) control is provided for comparison. One experiment using *mod-5(n3314)*, two experiments using *mod-5(n822)*, two experiments using the strain that restores MOD-5 expression in neurons including AIM and RIH (*AN::mod-5*), and two experiments using the strain that restores MOD-5 expression in neurons including NSM and ADF (*PN::mod-5*). Each circle represents one individual. Horizontal lines show means. In no experiment shown in this figure were hermaphrodites treated with ascr#10. Asterisks indicate levels of statistical significance (* for $p<0.05$, ** for $p<0.01$) for difference from the wild type using the Kolmogorov-Smirnov test.
